## Supplementary material for "Global, regional, and cryptic population structure in a high gene-flow transatlantic fish": Zip file containing all supplemental material.: Supplementary_Figures_Jansson_etal.docx

**Supplementary Figures for Jansson *et al*.**

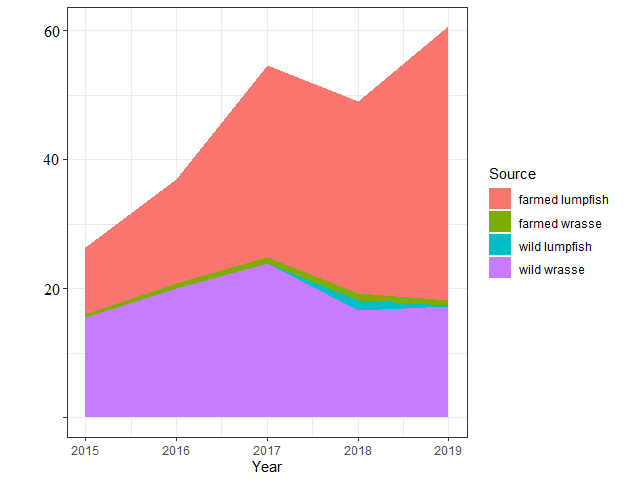

**S.Fig 1.** Use of cleaner fish in Norway in 2015-2019.

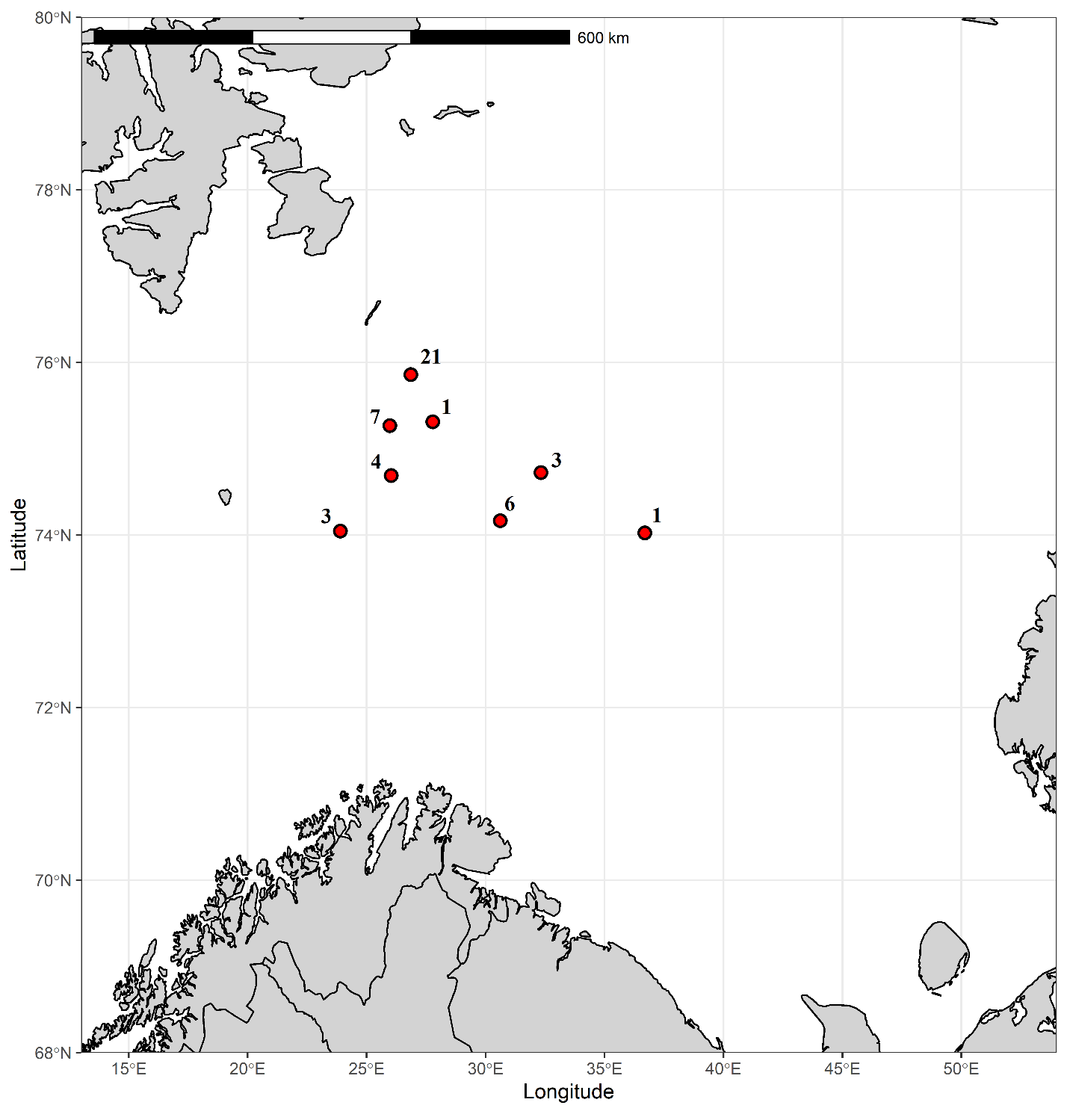

**S. Fig. 2a.** Samples collected from the Barents Sea (BAR_AR). Number besides each point denote number of collected from that exact location.

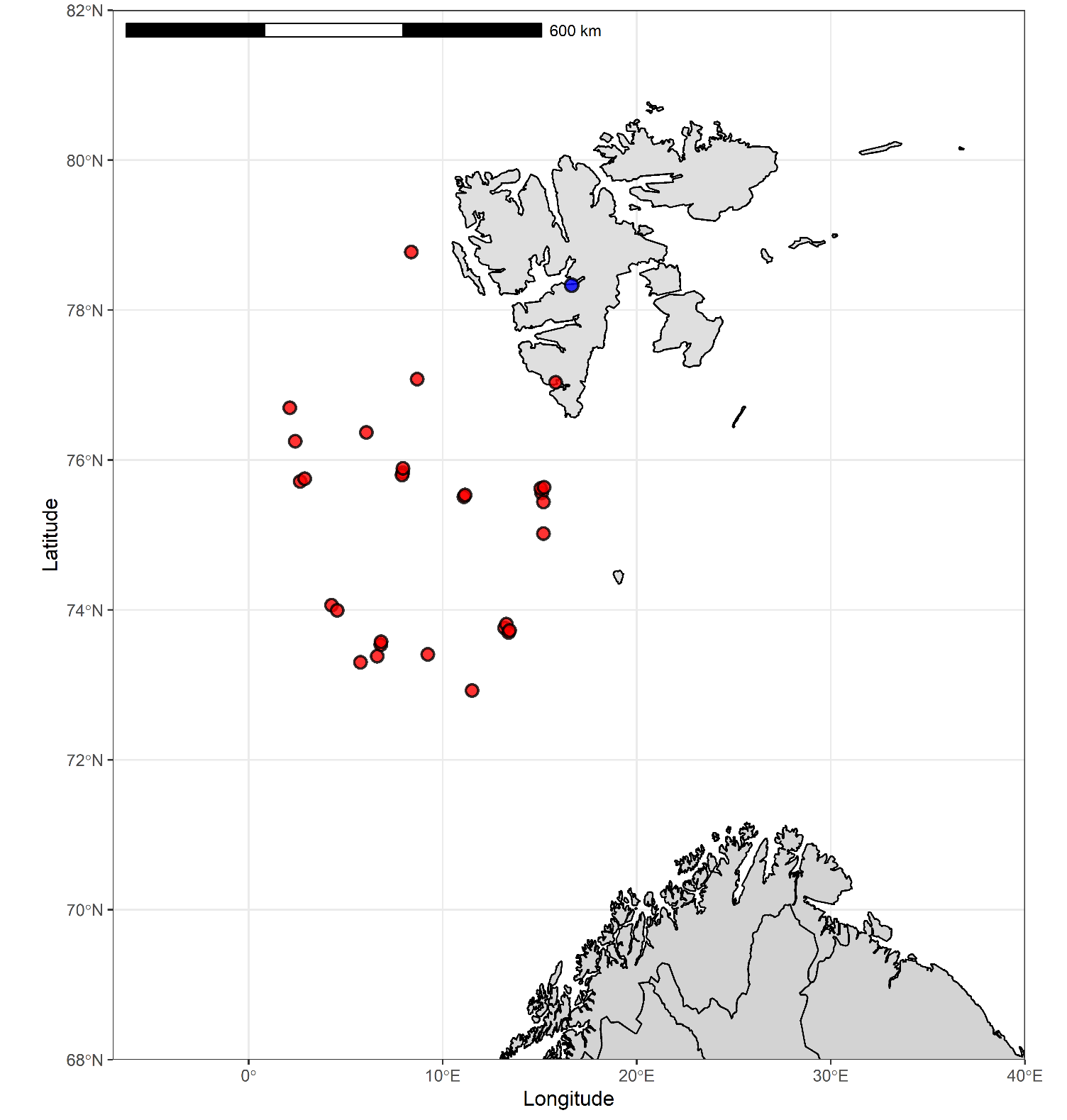

**S.Fig. 2b.** Samples collected close to Svalbard. Each red dot represent one fish (SVA_AR samples). Blue dot represent sampling point for all fish collected in Isfjorden (ISF_AR samples).

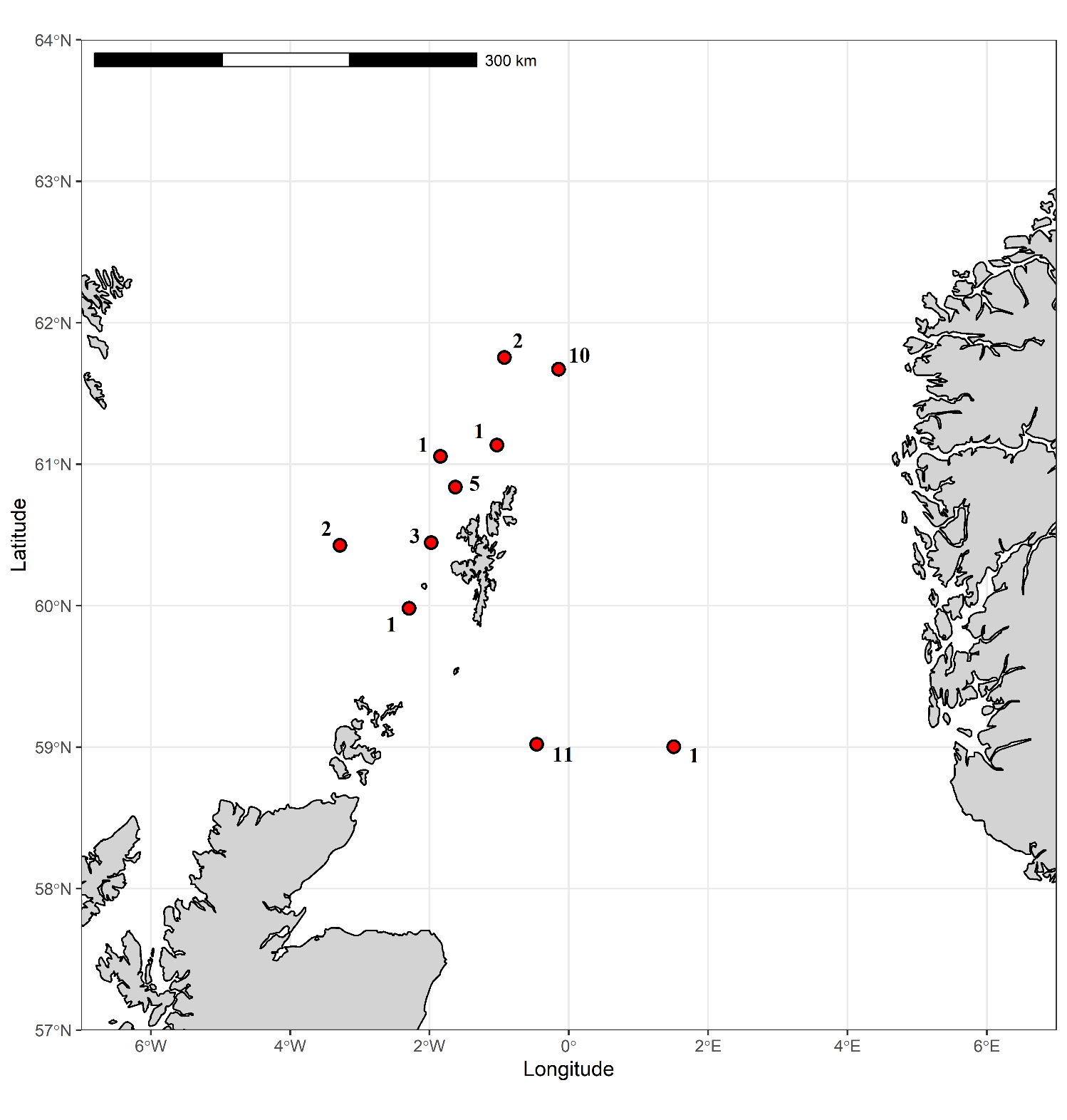

**S. Fig. 2c.** Samples from the North Sea. Red dots indicate sampling locations. Number besides each dot denotse number of fish collected from that exact location.

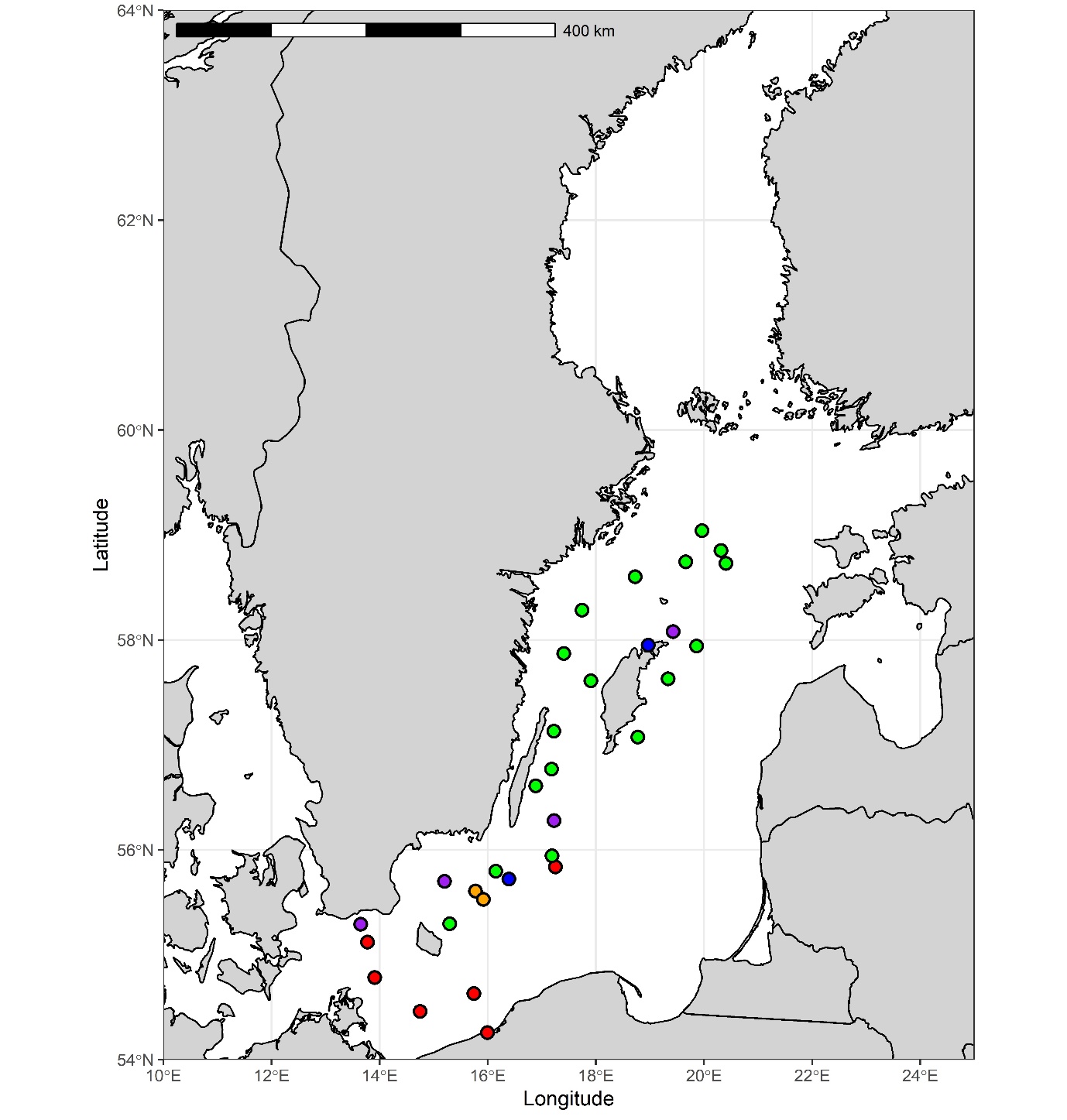

**S. Fig 2d.** Sampling locations for Baltic Sea samples. Different colors represent different sampling occations: BAL1 = red dots, BAL2 = blue dots, BAL3 = purple dots, BAL4 = green dots, BAL5 = orange dots (see Table 1). Note that several samples may be collected from same site.

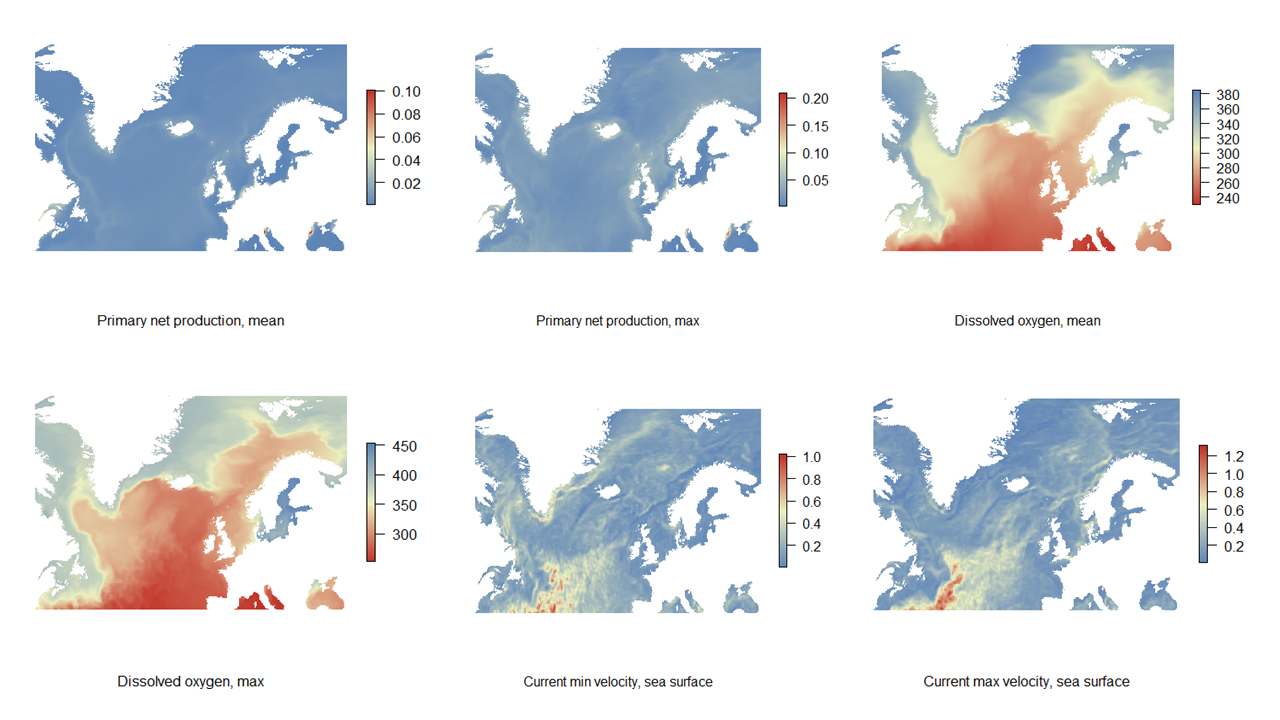

**S. Fig. 3a.** Visualization of the collected environmental data in the study area, first 6 parameters.

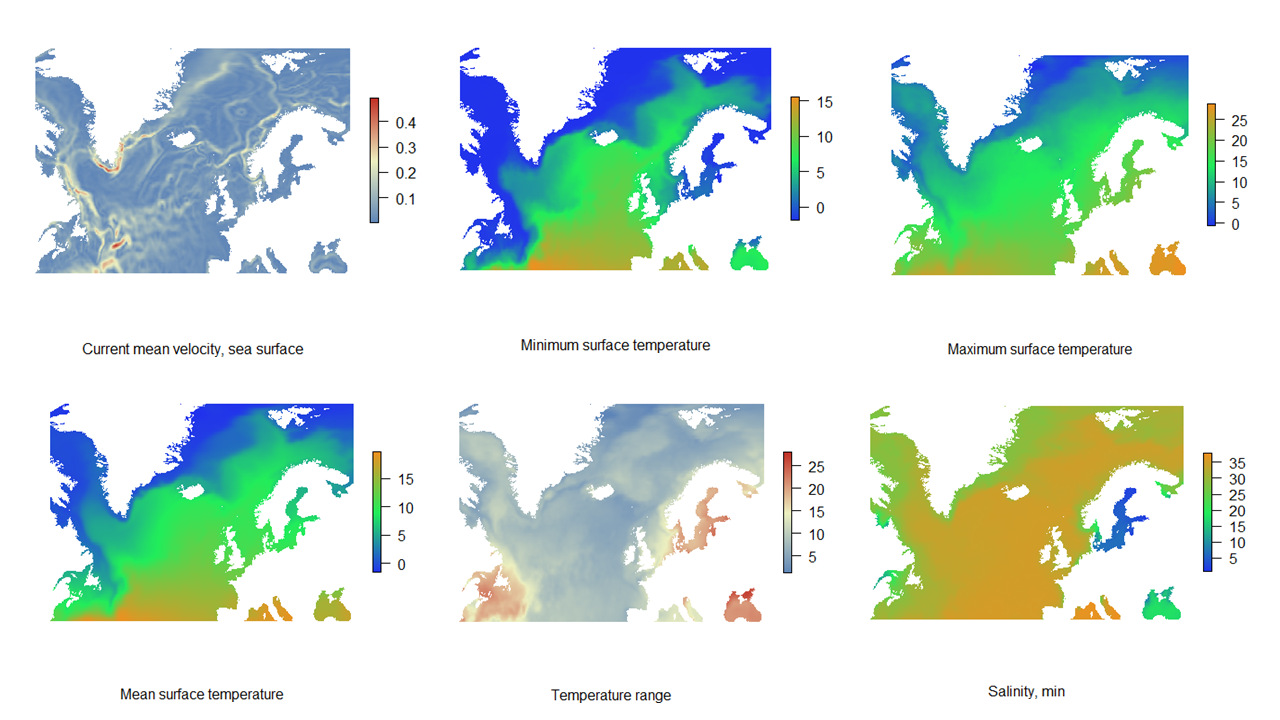

**S. Fig. 3b.** Visualization of the collected environmental data in the study area, next 6 parameters.

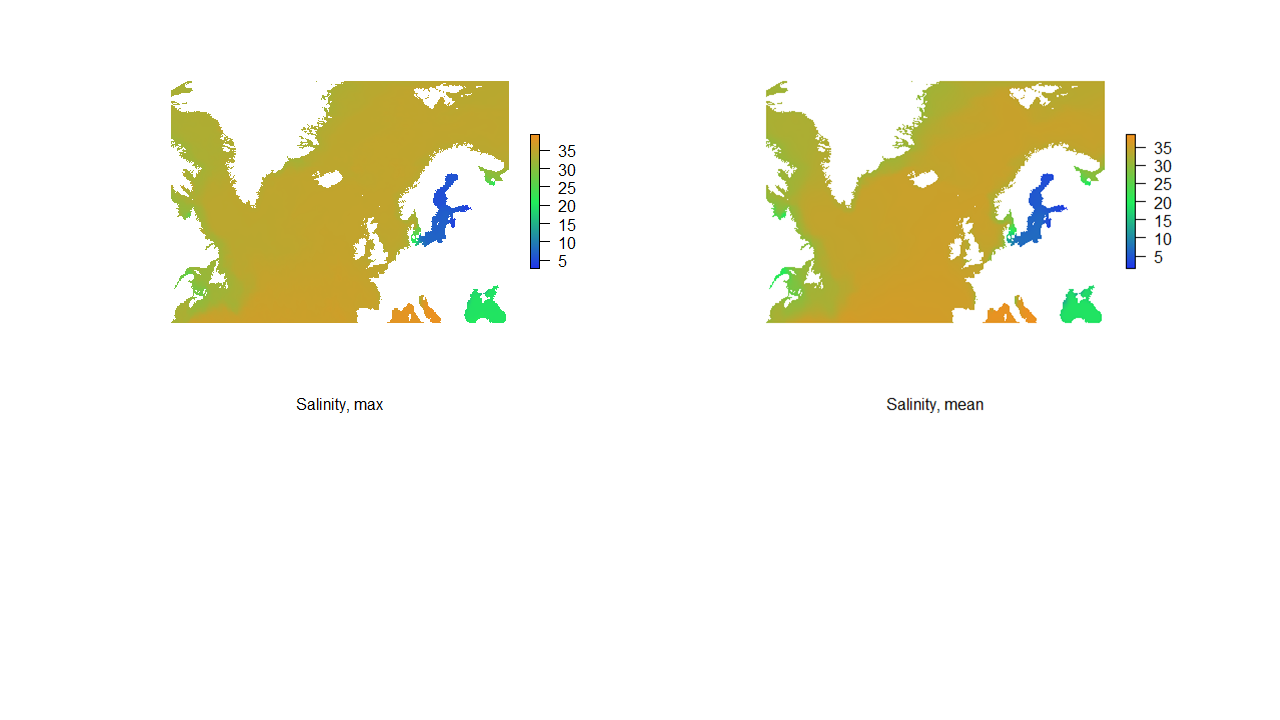

**S. Fig 3c.** Visualization of the collected environmental data in the study area, last two parameters.

**.**

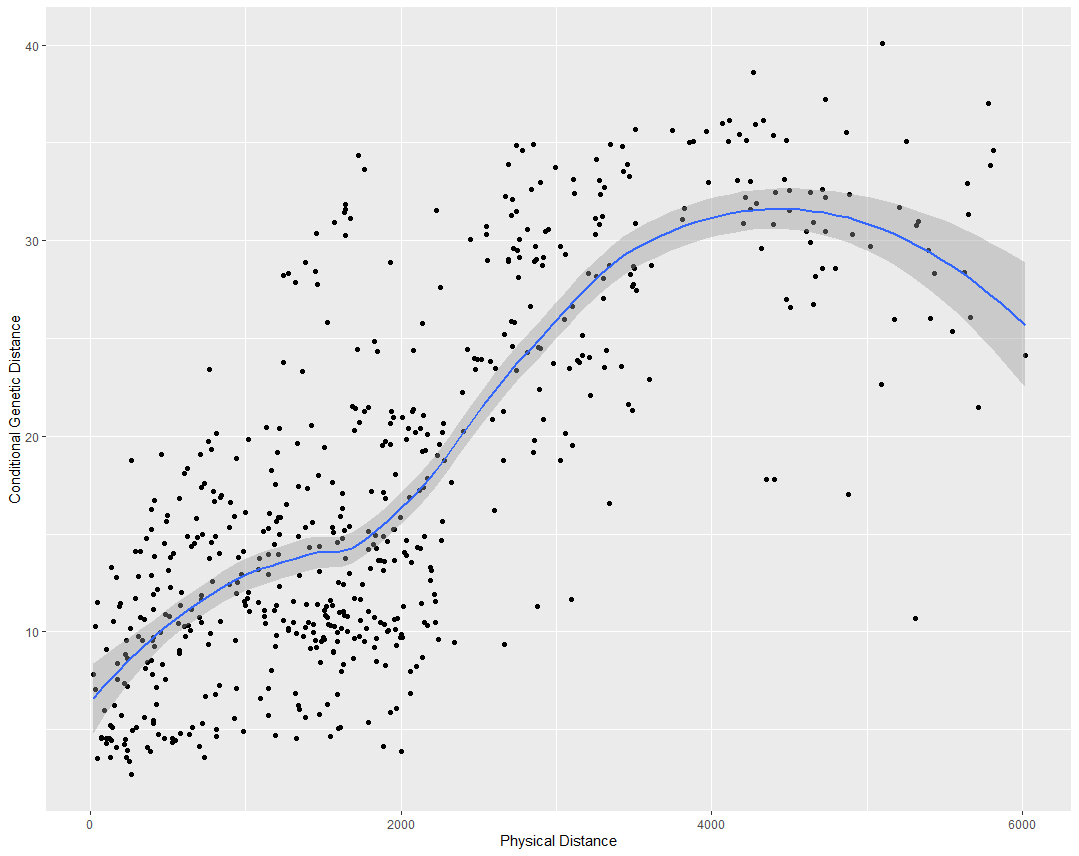

**S. Fig 4.a** Isolation by graph distance, IBGD

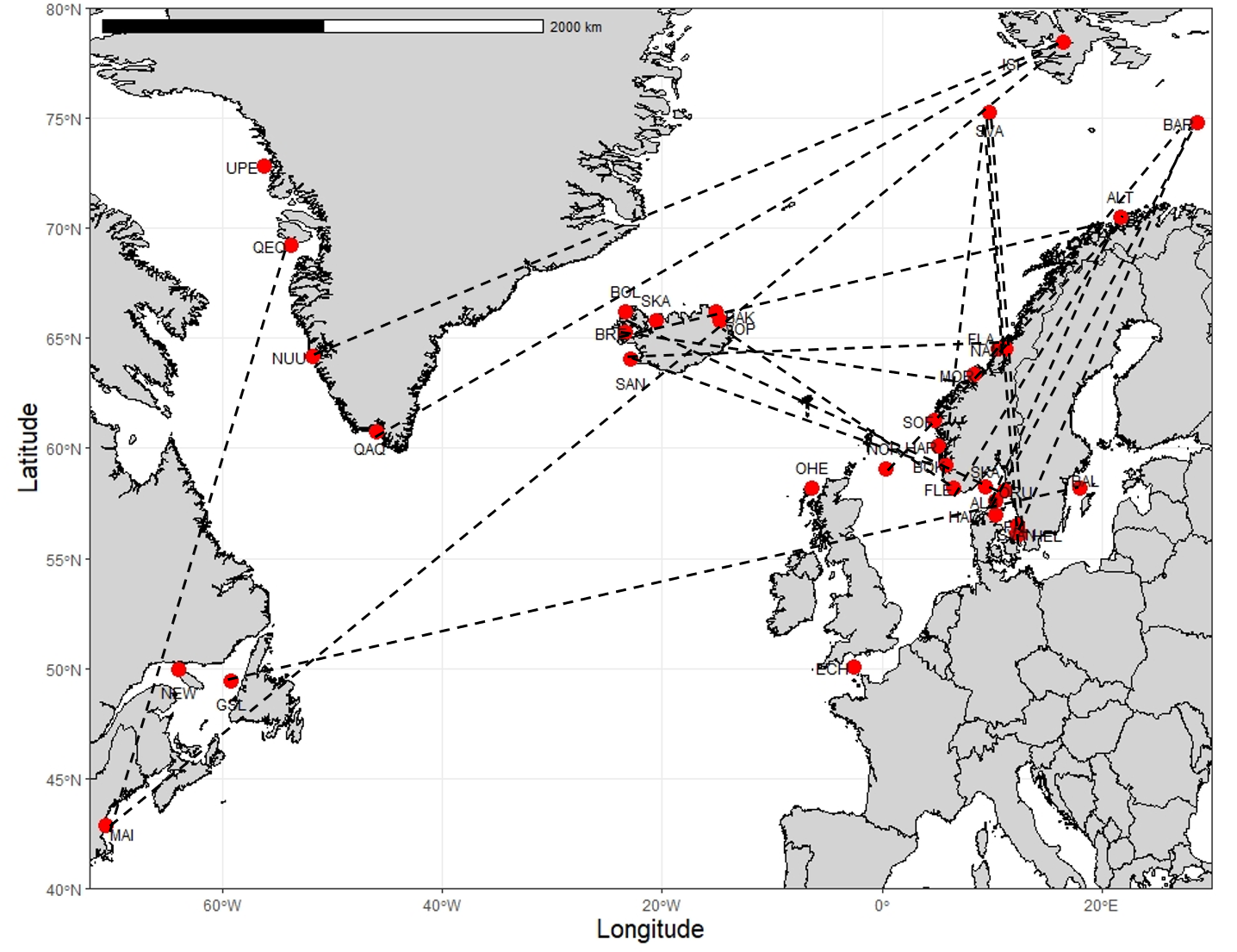

**S. Fig. 4b.** Map showing extended edges between sampling sites. Such edges are consistent with long-distance dispersal or other processes creating shared genetic diversity over simple IBD expectation.

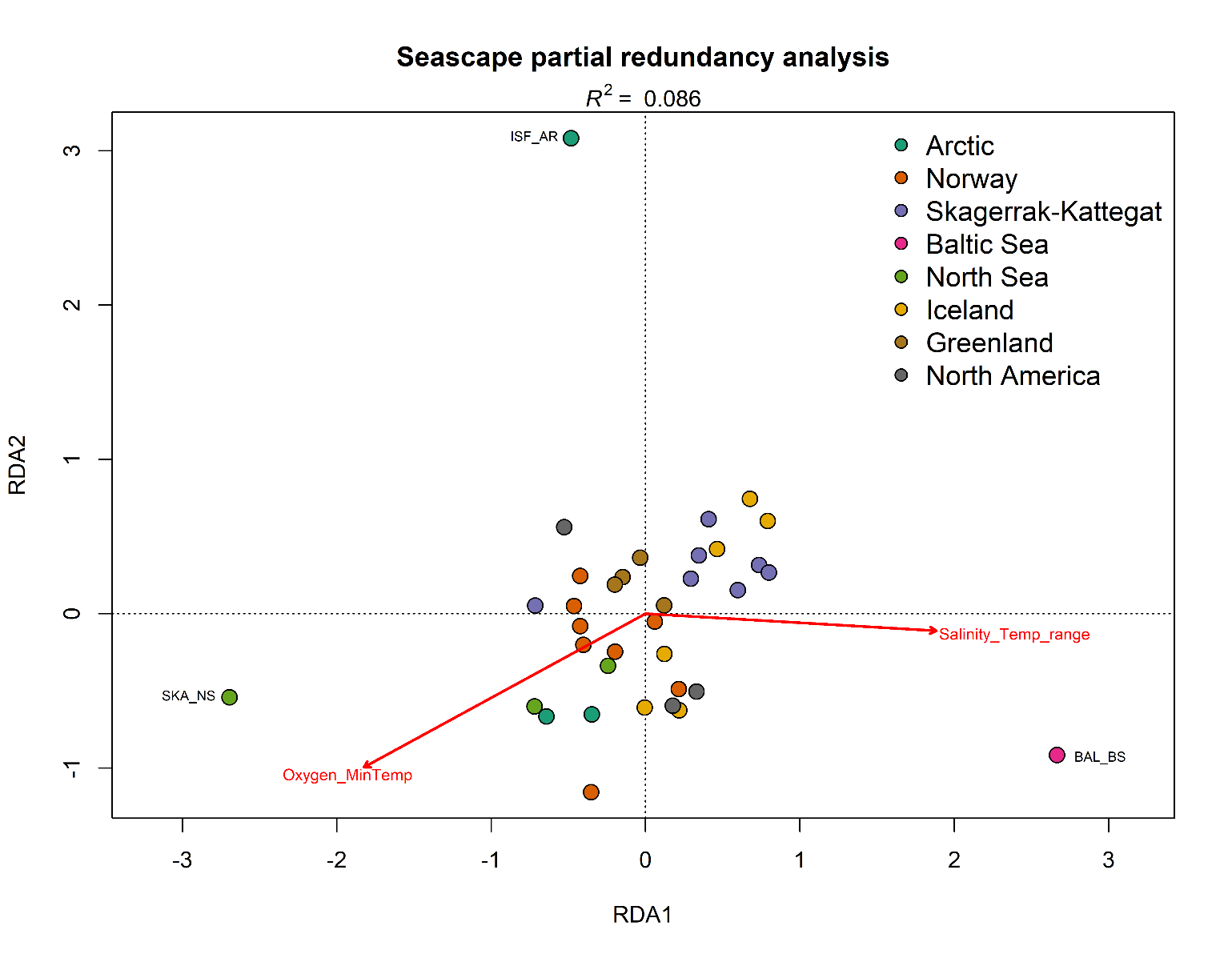

**S. Fig. 5.** Partial redundancy analysis between genetic and ecological matrices controlled by geographical location. Sampling locations are coloured based on their region.

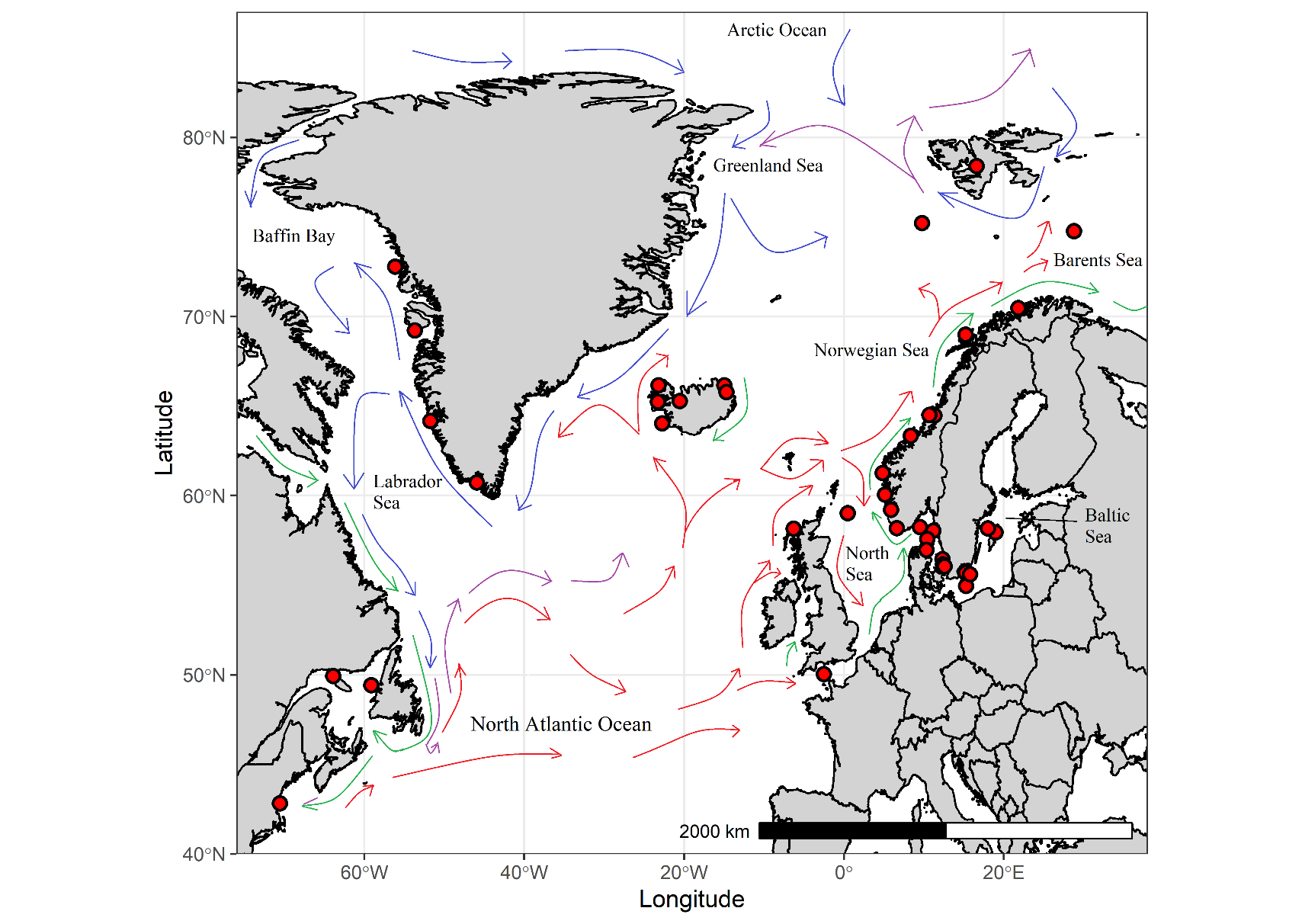

**S.Fig. 6.** Major surface currents in the study area. Sampling sites are shown with red circles. Green arrows indicate coastal currents, red arrows Atlantic currents, blue arrows Arctic currents, and purple arrows mixed Atlantic and Arctic currents. Note that the map is heavily distorted above ~77°. Source: <https://www.amap.no/documents/doc/major-surface-currents-in-the-north-atlantic-ocean/458> (sourced 25th of November, 2021).

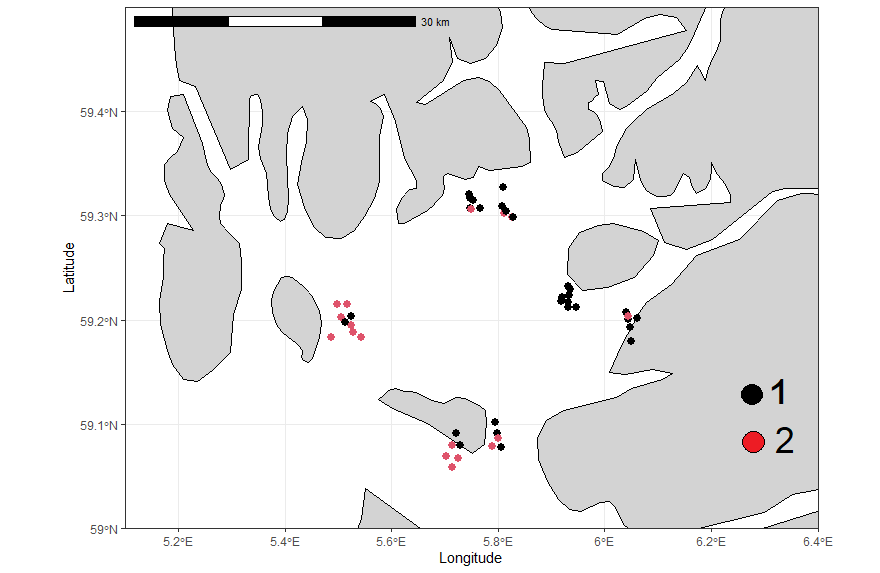

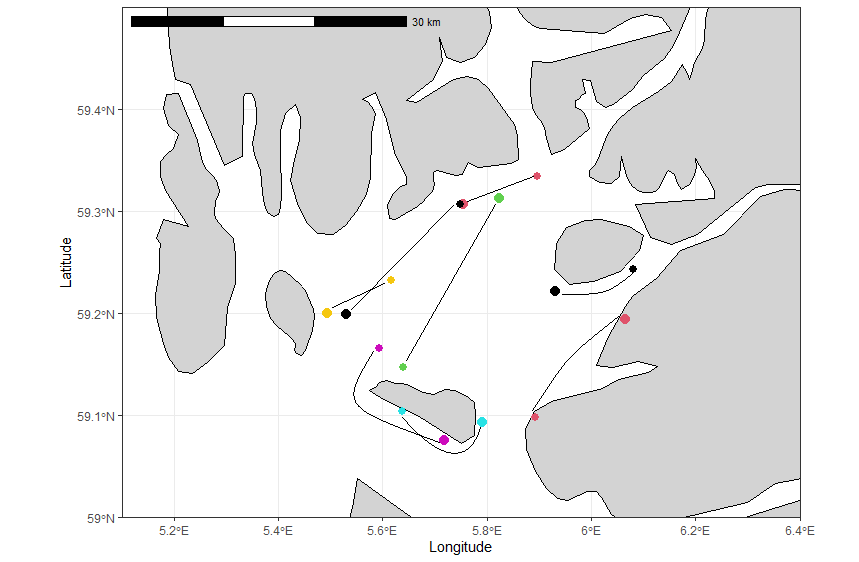

**S.Fig. 7.** Two genetic groups in the Boknafjord (BOK_NO). The figure above shows the likely genetic group membership of each caught fish in the trawl end position. The figure below shows approximate trawl routes along which fish were caught. Each colour represent different trawling days with smaller dots showing start positions and larger dots end position.
