## Supplementary material for "Global, regional, and cryptic population structure in a high gene-flow transatlantic fish": Zip file containing all supplemental material.: Supplementary_file_1_PCA_DAPC_plots_all.docx

**Supplementary file 1.** PCA and DAPC plots, all divisions as used in hierarchical clustering (Supplementary Table 5).

**The whole data, PCA (139 SNPs). 1597 fish from 39 locations.**

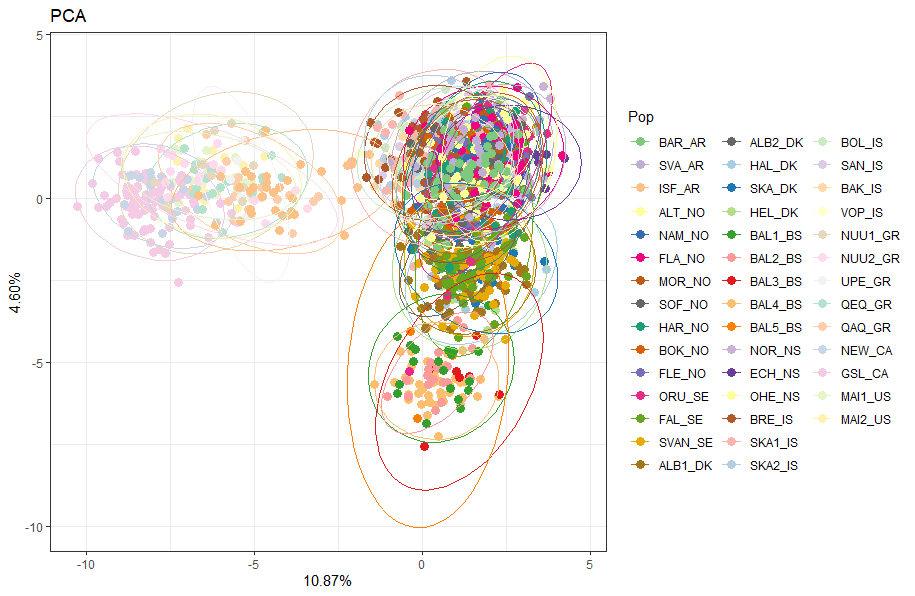

DAPC (130 PCs retained, 9 discriminant functions)

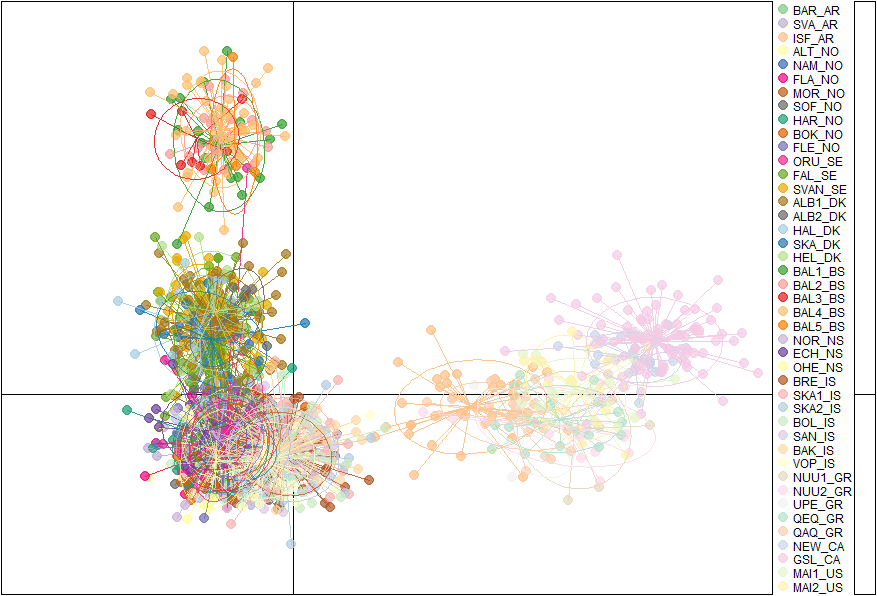

**The whole data, PCA (4393 SNPs). 95 fish from 10 locations.**

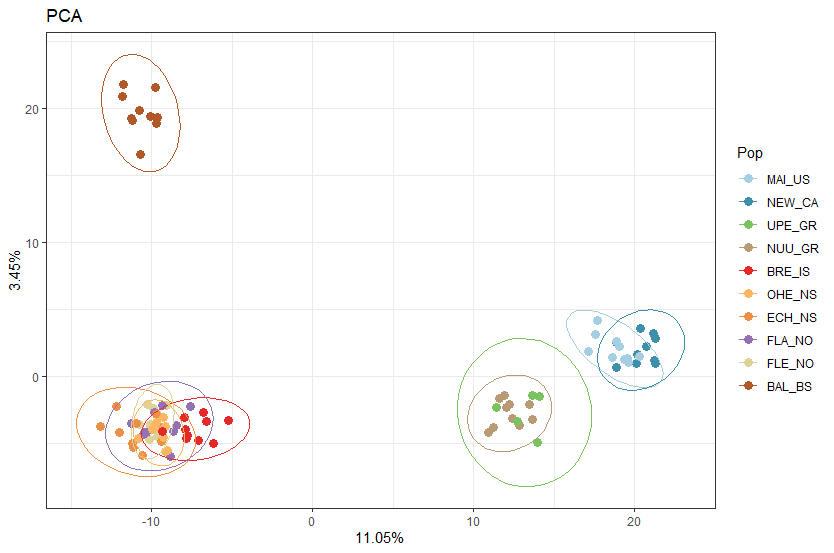

DAPC (90 PCs and 9 discriminant functions)

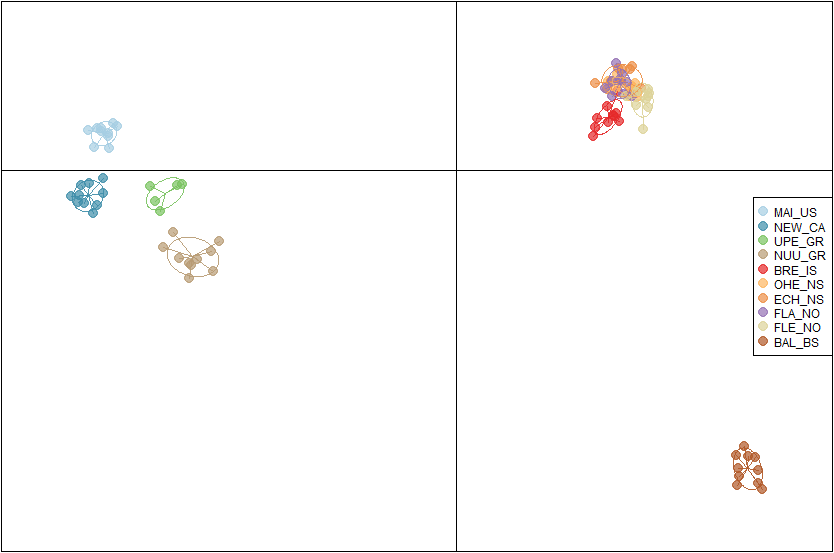

**Greenland and North America. (139 SNPs). 218 fish from 7 locations.**

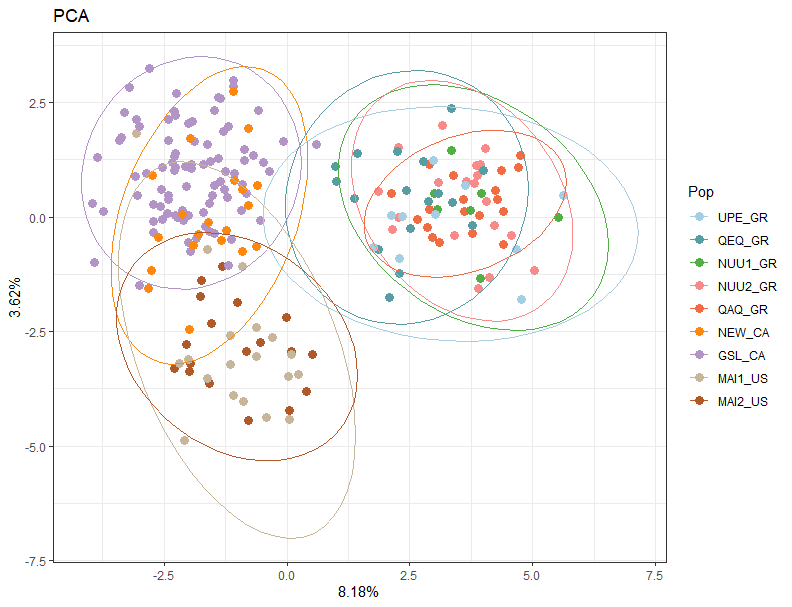

DAPC (60 PCs, 8 discriminant functions)

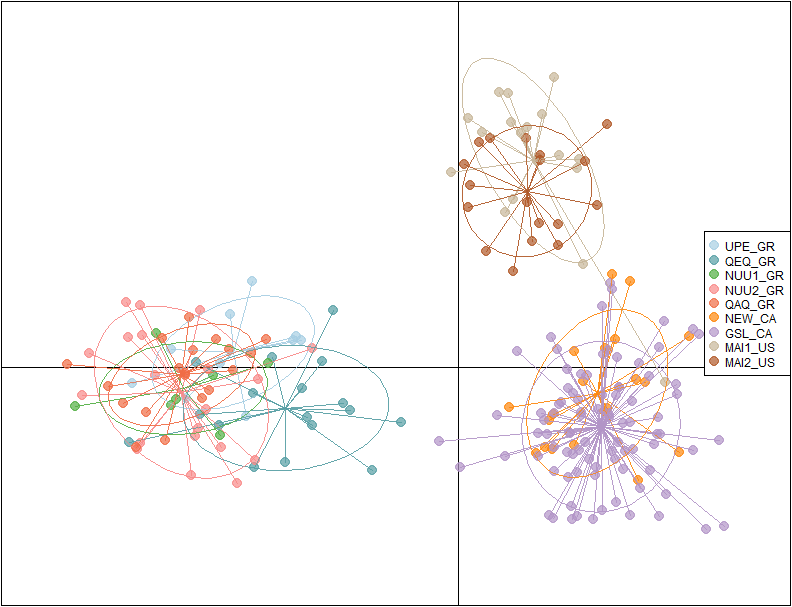

**PCA Iceland (139 SNPs). 386 fish from 6 locations.**

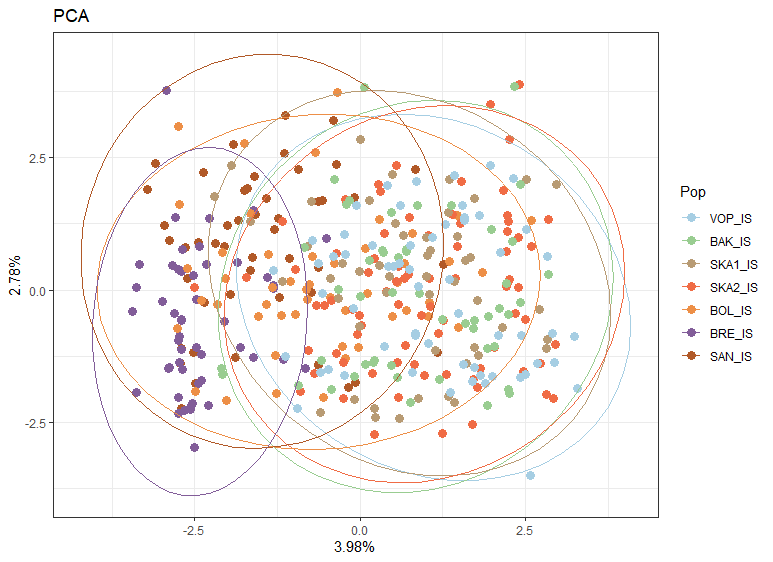

DAPC Iceland (80PCs, 6 discriminant functions)

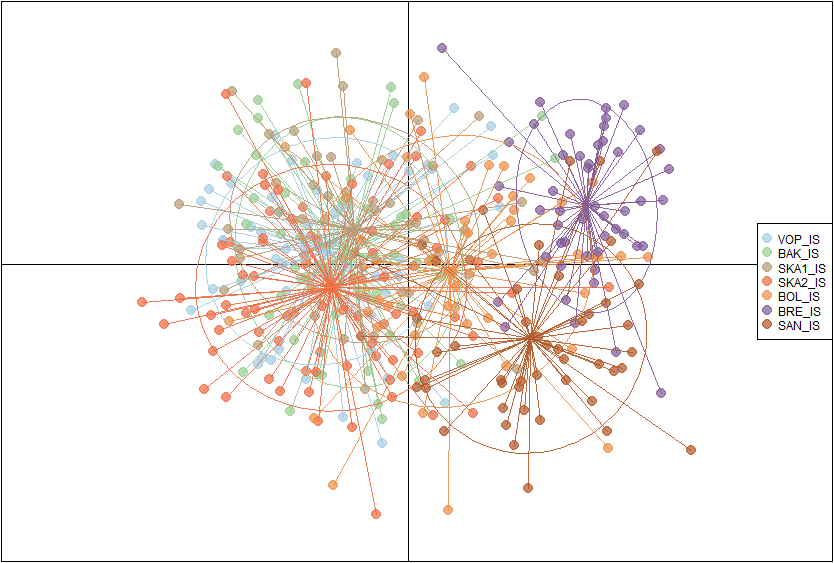

**Norway. (139 SNPs). 354 fish from 8 locations.**

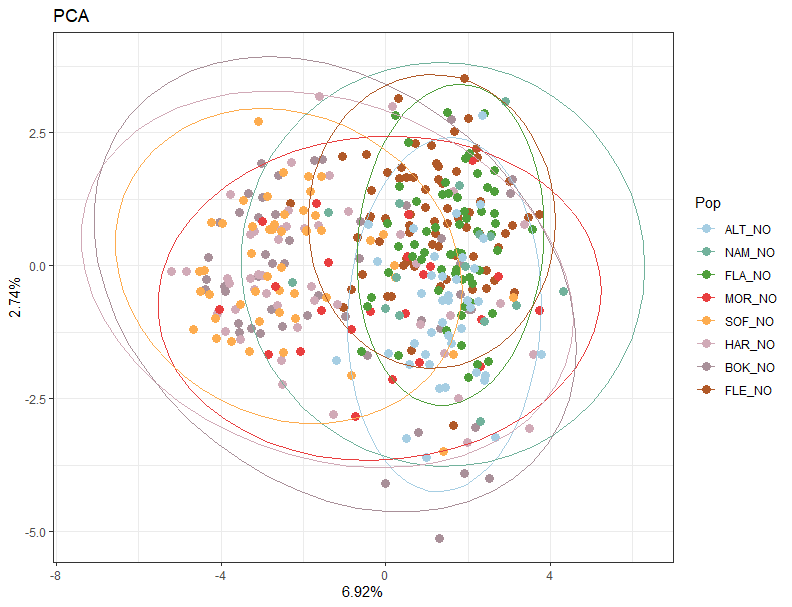

DAPC (80 PCs, 7 discriminant functions)

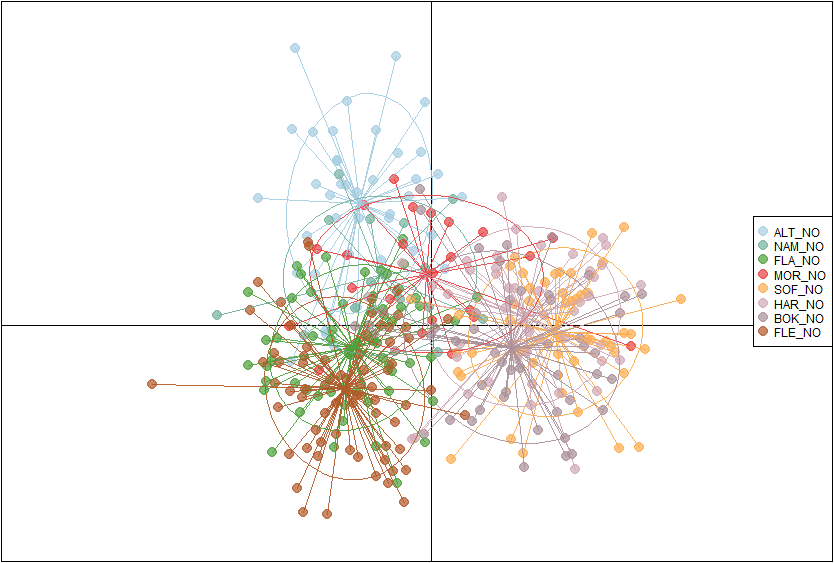

**Skagerrak, Kattegat, Baltic Sea. (139 SNPs). 451 fish from 12 locations.**

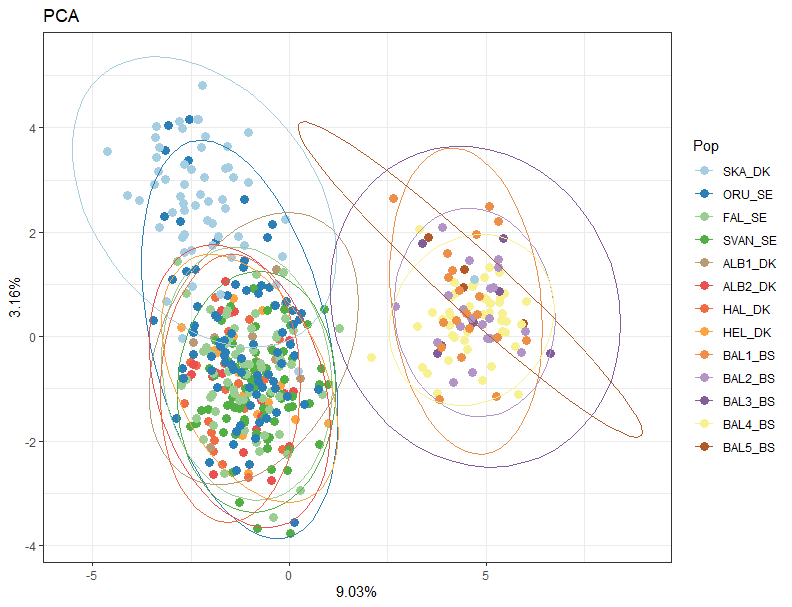

DAPC (120 PCs, 12 discriminant functions)

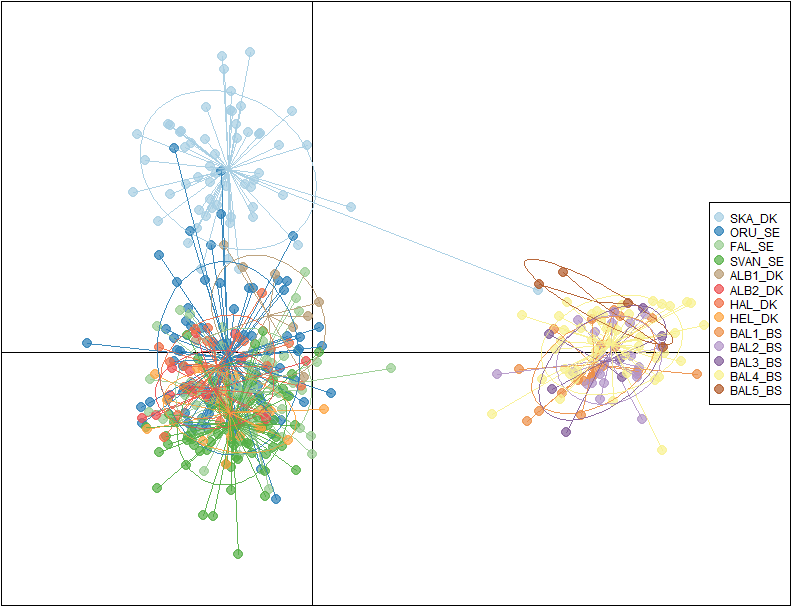

**Similar European populations with no clear assignment into any single population. (139 SNPs). 373 fish from 9 locations.**

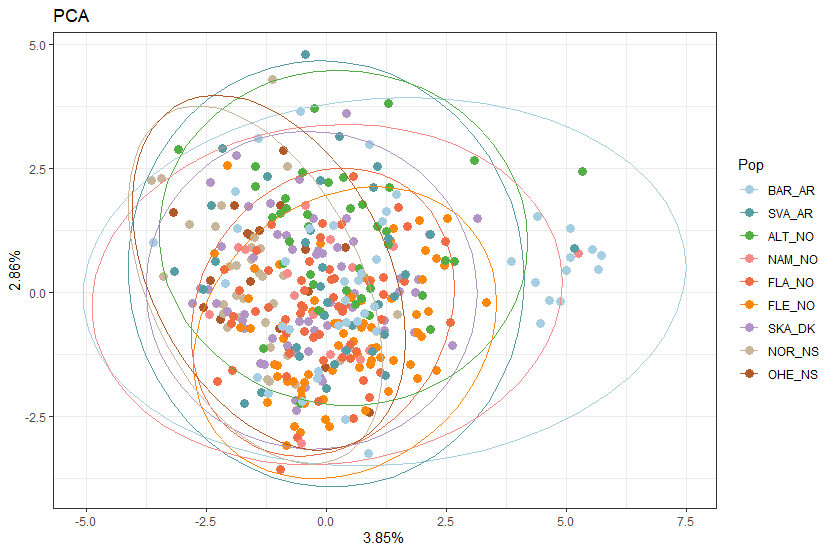

DAPC (50PCs, 8 discriminant functions)

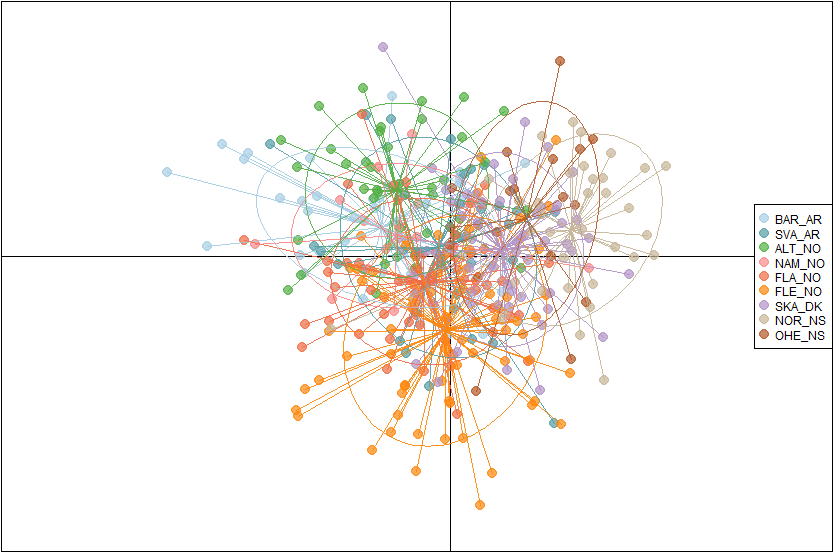

**Arctic sampling sites without Isfjorden (139 SNPs). 193 fish from 5 locations**

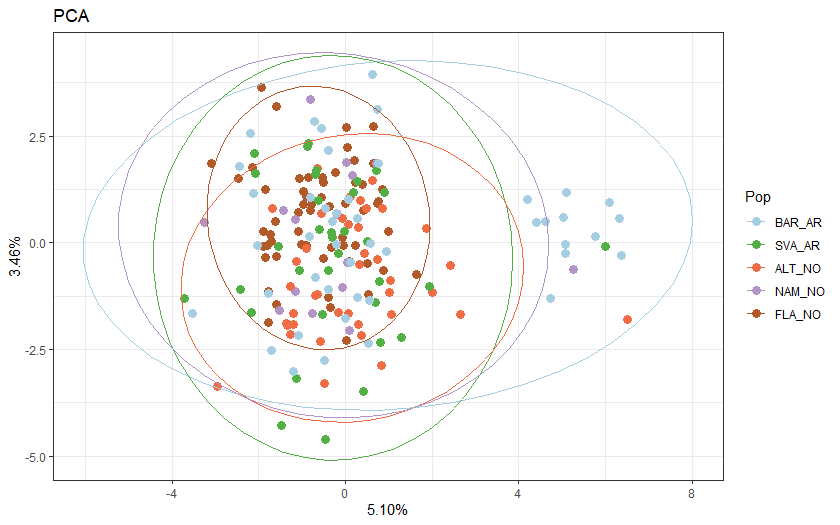

DAPC (40 PCs, 4 discriminant functions)

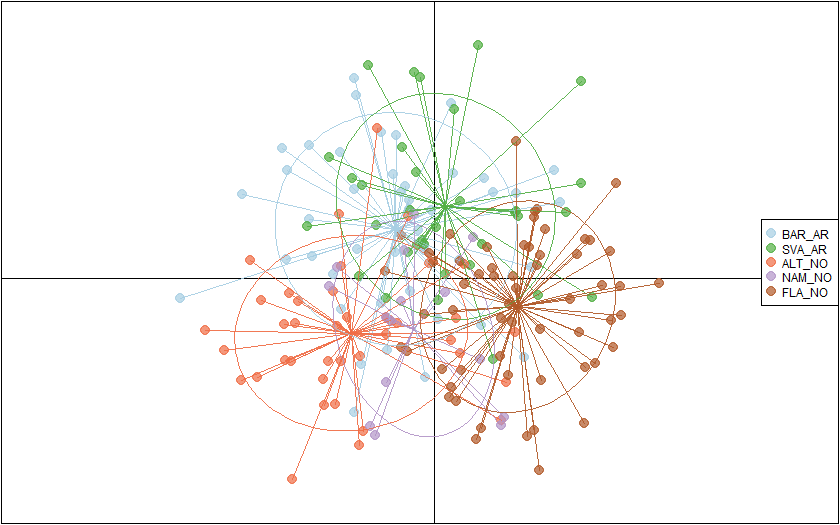
