## Supplementary material for "Global, regional, and cryptic population structure in a high gene-flow transatlantic fish": Zip file containing all supplemental material.: Supplementary_file_2_LFMM2_outliers.docx

**Supplementary File 2.** SNPs detected as outliers connected to environmental variables using *LFMM* method and reduced dataset of 139 SNPs. Below all analysis steps with results are shown.

**Step 1: Collinearity check.** If signficant correlation (-0.7≤ *R* ≥ 0.7) between a pair of variables was detected, variables in question were fused into same synthetic variable.
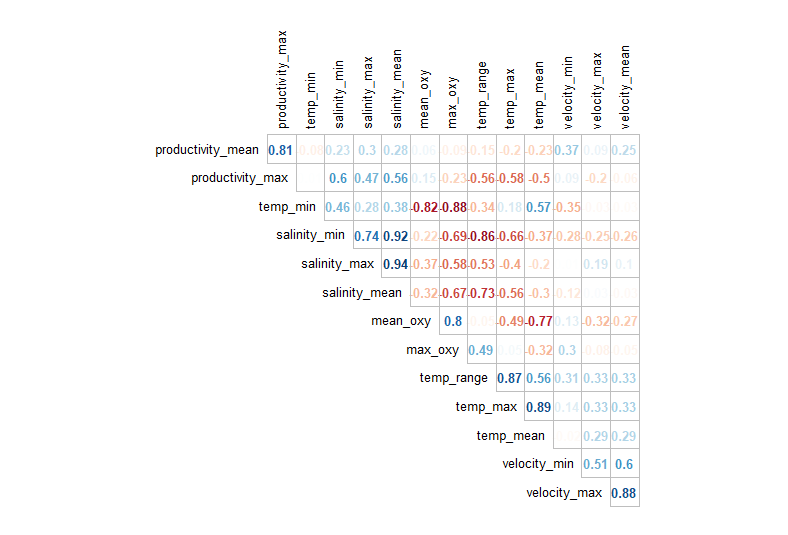

**Step 2: Regrouped correlated varibles, checked the stability of created synthetic (combined) variables (5) and re-checked correlation between them.**

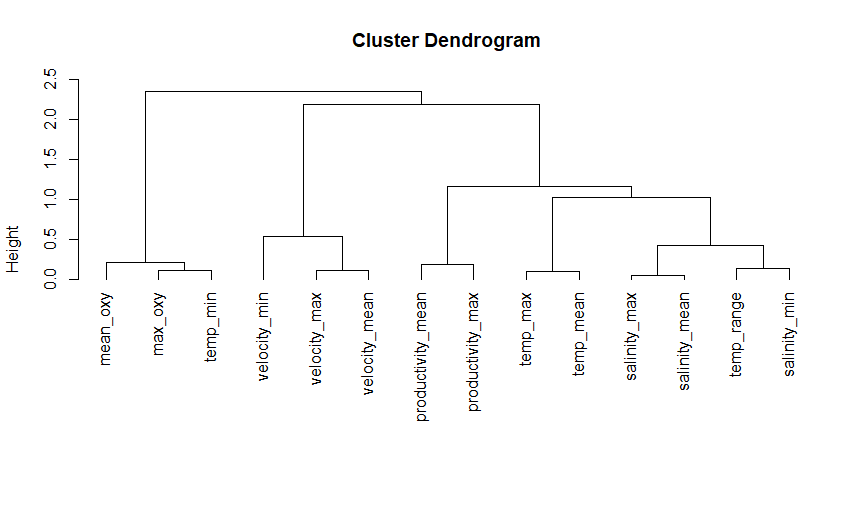

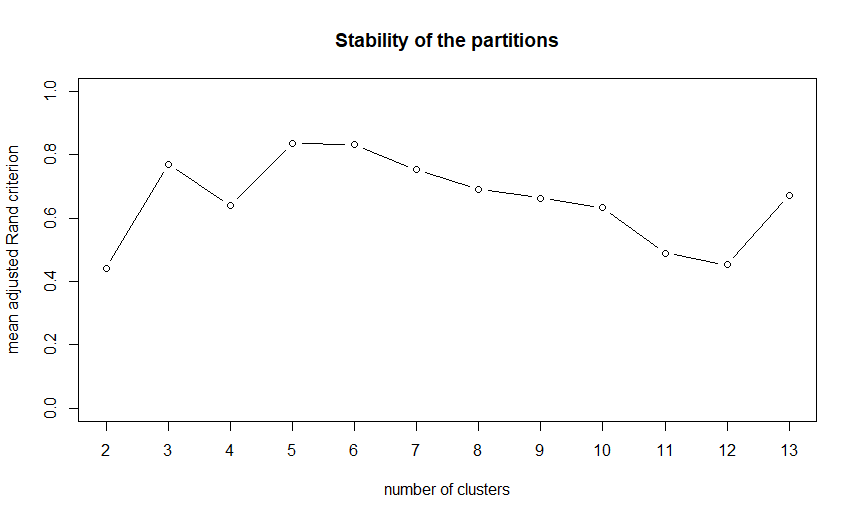

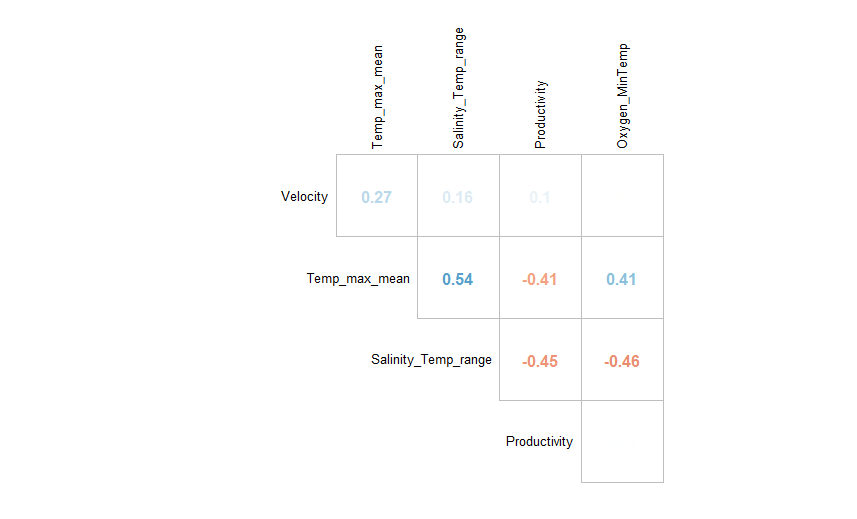

**Step 3. Ran LFMM analysis between these synthetic variables and imputed genetic dataset**. Based on preliminary PCA, *K* was set to 6. Checked distribution of p-values for all SNP-environment associations.

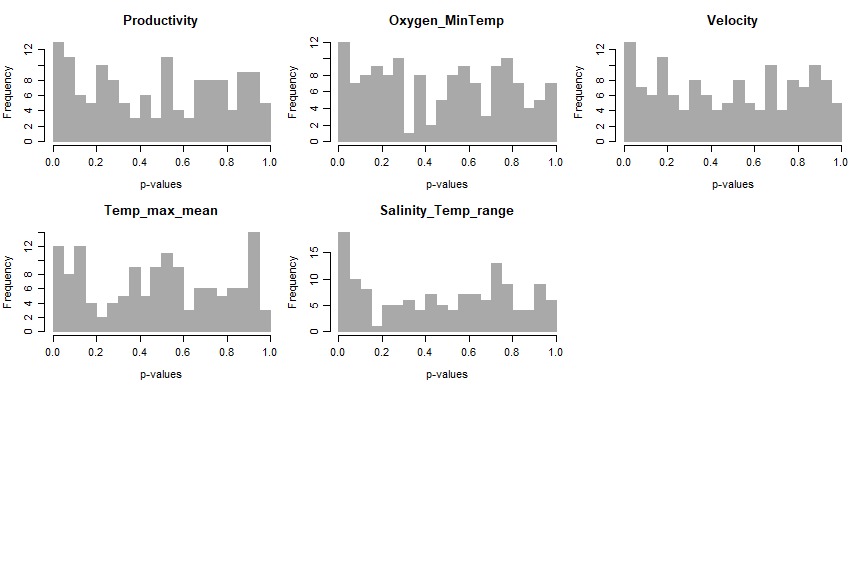

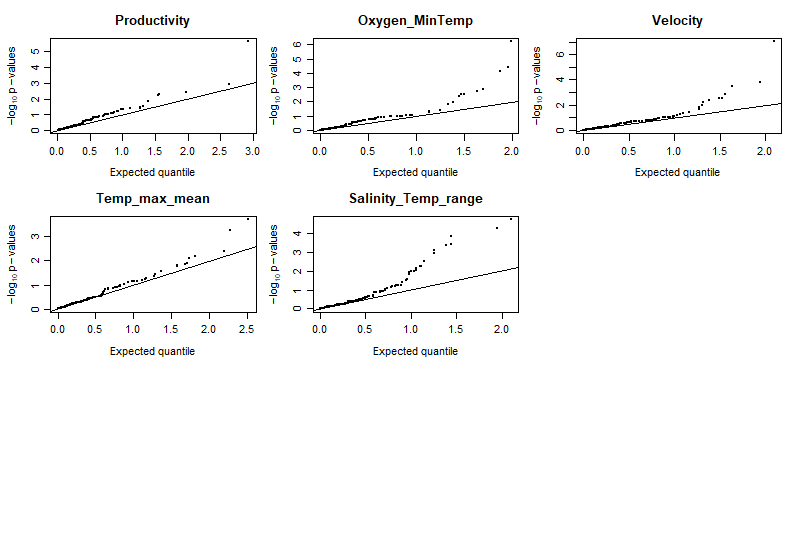

**Step 4. Derived q values from p values with false discovery rate (FDR) control.** Cut off of 5% was used. Below shown q value distribution for each synthetic variable and a table of significant SNPs using FDR of 5%. List of SNPs below the selected threshold is given as table below.

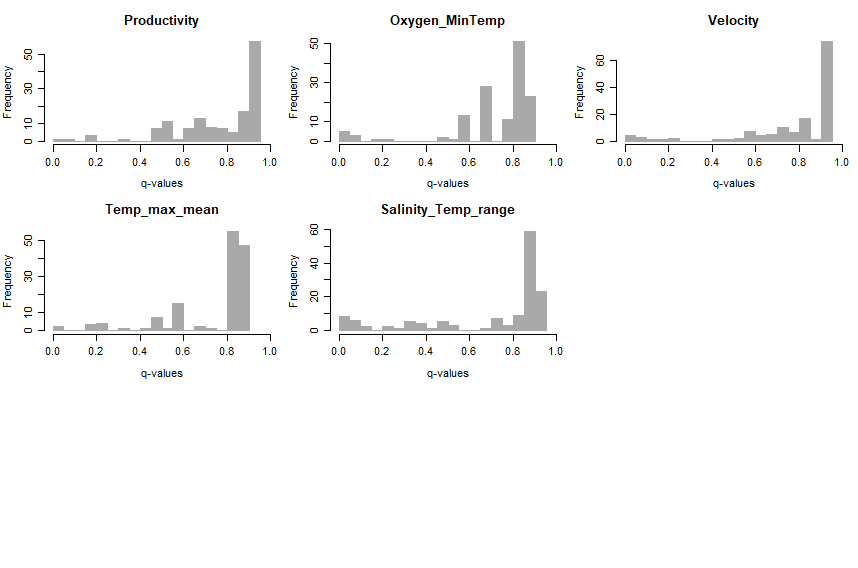

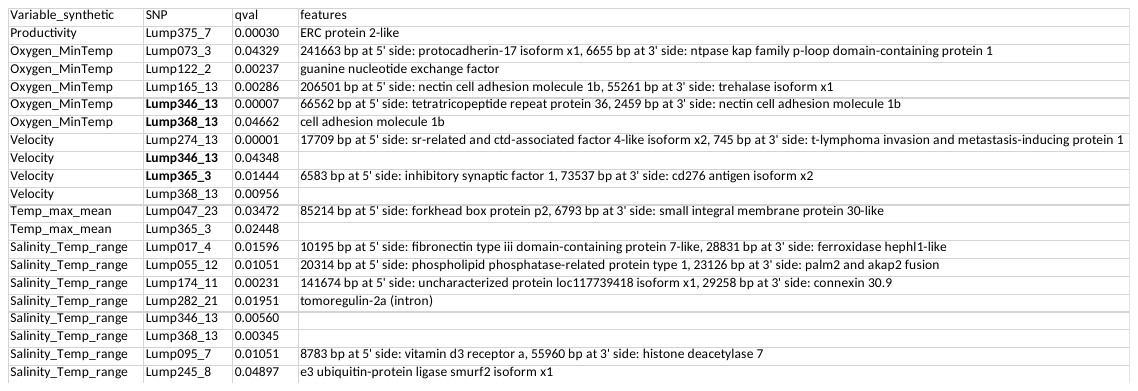

**Step 5. Visualize allelic distribution or the putatively selected loci divided into larger regions.** Populations are in same order as in Table 1 but VES_NO is removed, temporal replicates are combined and all Baltic Sea samples combined into one. Note that only one allele from each locus is shown.

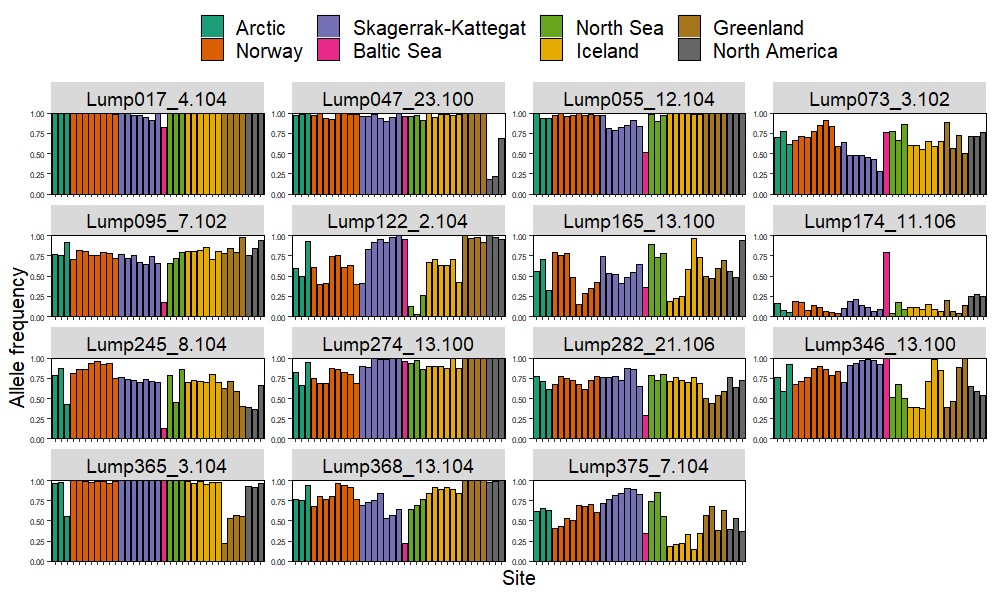
