## Supplementary material for "Global, regional, and cryptic population structure in a high gene-flow transatlantic fish": Zip file containing all supplemental material.: Supplementary_Table6_Popgraph_ types of edges.docx

**Table S6**. Popgraph analysis: The connections between nodes reflect the topology of genetic covariance amongst sampled site. Edges were classified based on the physical separation of sampling locations and the expectation given genetic covariance under a model of isolation by graph distance. When dispersal is uniform, the relative distances between nodes in Population Graph space should be proportional to physical (Euclidean) separation thus producing proportional edges. Extended edges indicate long-distance dispersal, *i.e.* sampling locations more spatially distant than expected by genetic covariance whereas compressed edges depict short distance dispersal (locations in closer proximity than expected by genetic covariance) suggesting barriers to gene flow.

| **Node1** | **Node2** | **Edge** | **Graph distance** | **Geographic distance (km)** | **Type of edge** |
| --- | --- | --- | --- | --- | --- |
| ECH_NS | UPE_GR | ECH_NS__UPE_GR | 22.89 | 3600 | Proportional |
| ISF_AR | UPE_GR | ISF_AR__UPE_GR | 13.64 | 1956 | Proportional |
| ECH_NS | NAM_NO | ECH_NS__NAM_NO | 13.27 | 1801 | Proportional |
| NEW_CA | NUU_GR | NEW_CA__NUU_GR | 11.68 | 1739 | Proportional |
| GSL_CA | NUU_GR | GSL_CA__NUU_GR | 11.70 | 1699 | Proportional |
| NEW_CA | QAQ_GR | NEW_CA__QAQ_GR | 13.78 | 1643 | Proportional |
| NAM_NO | OHE_NS | NAM_NO__OHE_NS | 8.00 | 1168 | Proportional |
| MAI_US | NEW_CA | MAI_US__NEW_CA | 7.09 | 942 | Proportional |
| NAM_NO | NOR_NS | NAM_NO__NOR_NS | 7.28 | 832 | Proportional |
| FLA_NO | NOR_NS | FLA_NO__NOR_NS | 5.02 | 812 | Proportional |
| ALT_NO | NAM_NO | ALT_NO__NAM_NO | 6.83 | 802 | Proportional |
| FLE_NO | NAM_NO | FLE_NO__NAM_NO | 6.68 | 743 | Proportional |
| FLA_NO | ORU_SE | FLA_NO__ORU_SE | 5.31 | 720 | Proportional |
| FLA_NO | SKA_DK | FLA_NO__SKA_DK | 4.15 | 699 | Proportional |
| ALT_NO | SVA_AR | ALT_NO__SVA_AR | 5.12 | 657 | Proportional |
| BOK_NO | FLA_NO | BOK_NO__FLA_NO | 4.76 | 640 | Proportional |
| FLE_NO | MOR_NO | FLE_NO__MOR_NO | 4.79 | 582 | Proportional |
| BAR_AR | SVA_AR | BAR_AR__SVA_AR | 4.48 | 548 | Proportional |
| NOR_NS | SKA_DK | NOR_NS__SKA_DK | 4.32 | 530 | Proportional |
| ALT_NO | BAR_AR | ALT_NO__BAR_AR | 4.57 | 529 | Proportional |
| BOK_NO | MOR_NO | BOK_NO__MOR_NO | 4.57 | 479 | Proportional |
| BAK_IS | SAN_IS | BAK_IS__SAN_IS | 4.75 | 435 | Proportional |
| BOL_IS | VOP_IS | BOL_IS__VOP_IS | 3.89 | 388 | Proportional |
| BAK_IS | BOL_IS | BAK_IS__BOL_IS | 4.11 | 370 | Proportional |
| SKA_IS | VOP_IS | SKA_IS__VOP_IS | 2.69 | 263 | Proportional |
| BAK_IS | SKA_IS | BAK_IS__SKA_IS | 3.39 | 252 | Proportional |
| BOK_NO | SOF_NO | BOK_NO__SOF_NO | 3.57 | 235 | Proportional |
| ISF_AR | MAI_US | ISF_AR__MAI_US | 10.70 | 5308 | Extended |
| BAL_BS | GSL_CA | BAL_BS__GSL_CA | 17.04 | 4877 | Extended |
| BAL_BS | UPE_GR | BAL_BS__UPE_GR | 21.30 | 3487 | Extended |
| FAL_SE | UPE_GR | FAL_SE__UPE_GR | 21.63 | 3463 | Extended |
| HEL_DK | QAQ_GR | HEL_DK__QAQ_GR | 16.54 | 3338 | Extended |
| MAI_US | QEQ_GR | MAI_US__QEQ_GR | 11.67 | 3094 | Extended |
| ISF_AR | QAQ_GR | ISF_AR__QAQ_GR | 11.30 | 2879 | Extended |
| ISF_AR | NUU_GR | ISF_AR__NUU_GR | 9.37 | 2664 | Extended |
| GSL_CA | QEQ_GR | GSL_CA__QEQ_GR | 11.54 | 2223 | Extended |
| ISF_AR | QEQ_GR | ISF_AR__QEQ_GR | 10.50 | 2222 | Extended |
| NEW_CA | QEQ_GR | NEW_CA__QEQ_GR | 11.89 | 2215 | Extended |
| HEL_DK | SVA_AR | HEL_DK__SVA_AR | 8.68 | 2136 | Extended |
| ALB_DK | BAR_AR | ALB_DK__BAR_AR | 6.88 | 2061 | Extended |
| BAR_AR | SKA_DK | BAR_AR__SKA_DK | 3.89 | 2005 | Extended |
| BOL_IS | ORU_SE | BOL_IS__ORU_SE | 6.09 | 1972 | Extended |
| ALT_NO | BRE_IS | ALT_NO__BRE_IS | 5.90 | 1928 | Extended |
| SKA_DK | SVA_AR | SKA_DK__SVA_AR | 4.16 | 1889 | Extended |
| BOK_NO | SVA_AR | BOK_NO__SVA_AR | 5.36 | 1788 | Extended |
| BRE_IS | NAM_NO | BRE_IS__NAM_NO | 7.98 | 1617 | Extended |
| ALT_NO | NOR_NS | ALT_NO__NOR_NS | 5.13 | 1611 | Extended |
| FLA_NO | SAN_IS | FLA_NO__SAN_IS | 5.06 | 1599 | Extended |
| BOK_NO | SAN_IS | BOK_NO__SAN_IS | 6.81 | 1591 | Extended |
| ALT_NO | FLE_NO | ALT_NO__FLE_NO | 4.67 | 1542 | Extended |
| BRE_IS | MOR_NO | BRE_IS__MOR_NO | 6.27 | 1526 | Extended |
| ALT_NO | ORU_SE | ALT_NO__ORU_SE | 5.79 | 1471 | Extended |
| FLE_NO | VOP_IS | FLE_NO__VOP_IS | 5.63 | 1387 | Extended |
| NOR_NS | SAN_IS | NOR_NS__SAN_IS | 6.03 | 1344 | Extended |
| BAK_IS | SVA_AR | BAK_IS__SVA_AR | 6.24 | 1338 | Extended |
| BAR_AR | FLA_NO | BAR_AR__FLA_NO | 4.54 | 1327 | Extended |
| BAR_AR | NAM_NO | BAR_AR__NAM_NO | 6.86 | 1318 | Extended |
| FLA_NO | SVA_AR | FLA_NO__SVA_AR | 4.73 | 1193 | Extended |
| GSL_CA | MAI_US | GSL_CA__MAI_US | 7.11 | 1145 | Extended |
| FLA_NO | OHE_NS | FLA_NO__OHE_NS | 5.74 | 1144 | Extended |
| OHE_NS | SAN_IS | OHE_NS__SAN_IS | 6.62 | 1096 | Extended |
| ALT_NO | MOR_NO | ALT_NO__MOR_NO | 4.93 | 982 | Extended |
| OHE_NS | SKA_DK | OHE_NS__SKA_DK | 5.57 | 926 | Extended |
| ALT_NO | FLA_NO | ALT_NO__FLA_NO | 4.63 | 814 | Extended |
| FLA_NO | FLE_NO | FLA_NO__FLE_NO | 3.60 | 736 | Extended |
| BAL_BS | ECH_NS | BAL_BS__ECH_NS | 15.92 | 1607 | Compressed |
| QAQ_GR | UPE_GR | QAQ_GR__UPE_GR | 14.32 | 1410 | Compressed |
| BRE_IS | NUU_GR | BRE_IS__NUU_GR | 12.89 | 1348 | Compressed |
| ECH_NS | FAL_SE | ECH_NS__FAL_SE | 14.96 | 1215 | Compressed |
| ECH_NS | HEL_DK | ECH_NS__HEL_DK | 13.94 | 1209 | Compressed |
| ECH_NS | HAL_DK | ECH_NS__HAL_DK | 12.96 | 1145 | Compressed |
| ECH_NS | FLE_NO | ECH_NS__FLE_NO | 11.52 | 1083 | Compressed |
| ECH_NS | NOR_NS | ECH_NS__NOR_NS | 12.03 | 1018 | Compressed |
| QAQ_GR | QEQ_GR | QAQ_GR__QEQ_GR | 11.71 | 1010 | Compressed |
| NUU_GR | UPE_GR | NUU_GR__UPE_GR | 12.93 | 975 | Compressed |
| ECH_NS | OHE_NS | ECH_NS__OHE_NS | 11.98 | 937 | Compressed |
| FAL_SE | NAM_NO | FAL_SE__NAM_NO | 12.43 | 895 | Compressed |
| BAL_BS | NAM_NO | BAL_BS__NAM_NO | 12.57 | 787 | Compressed |
| FAL_SE | HAR_NO | FAL_SE__HAR_NO | 11.35 | 579 | Compressed |
| NUU_GR | QEQ_GR | NUU_GR__QEQ_GR | 10.43 | 568 | Compressed |
| BAR_AR | ISF_AR | BAR_AR__ISF_AR | 10.79 | 510 | Compressed |
| NUU_GR | QAQ_GR | NUU_GR__QAQ_GR | 10.90 | 485 | Compressed |
| NAM_NO | SOF_NO | NAM_NO__SOF_NO | 7.59 | 485 | Compressed |
| ALB_DK | BAL_BS | ALB_DK__BAL_BS | 9.97 | 453 | Compressed |
| SKA_DK | SOF_NO | SKA_DK__SOF_NO | 6.31 | 425 | Compressed |
| QEQ_GR | UPE_GR | QEQ_GR__UPE_GR | 11.91 | 407 | Compressed |
| BAL_BS | SVAN_SE | BAL_BS__SVAN_SE | 9.71 | 407 | Compressed |
| NOR_NS | OHE_NS | NOR_NS__OHE_NS | 5.47 | 404 | Compressed |
| HAR_NO | MOR_NO | HAR_NO__MOR_NO | 5.32 | 403 | Compressed |
| BAL_BS | HEL_DK | BAL_BS__HEL_DK | 11.14 | 403 | Compressed |
| BAL_BS | ORU_SE | BAL_BS__ORU_SE | 9.55 | 401 | Compressed |
| BAL_BS | FAL_SE | BAL_BS__FAL_SE | 12.89 | 392 | Compressed |
| GSL_CA | NEW_CA | GSL_CA__NEW_CA | 5.63 | 347 | Compressed |
| MOR_NO | SOF_NO | MOR_NO__SOF_NO | 5.13 | 295 | Compressed |
| FLE_NO | ORU_SE | FLE_NO__ORU_SE | 4.98 | 269 | Compressed |
| BOL_IS | SAN_IS | BOL_IS__SAN_IS | 3.96 | 237 | Compressed |
| HEL_DK | ORU_SE | HEL_DK__ORU_SE | 7.21 | 237 | Compressed |
| SAN_IS | SKA_IS | SAN_IS__SKA_IS | 4.48 | 223 | Compressed |
| ORU_SE | SVAN_SE | ORU_SE__SVAN_SE | 4.25 | 222 | Compressed |
| ALB_DK | HEL_DK | ALB_DK__HEL_DK | 7.34 | 216 | Compressed |
| ALB_DK | SVAN_SE | ALB_DK__SVAN_SE | 5.74 | 199 | Compressed |
| HAL_DK | HEL_DK | HAL_DK__HEL_DK | 7.56 | 172 | Compressed |
| FLE_NO | SKA_DK | FLE_NO__SKA_DK | 4.11 | 170 | Compressed |
| HAL_DK | SVAN_SE | HAL_DK__SVAN_SE | 6.23 | 155 | Compressed |
| BRE_IS | SKA_IS | BRE_IS__SKA_IS | 5.13 | 144 | Compressed |
| BRE_IS | SAN_IS | BRE_IS__SAN_IS | 4.46 | 136 | Compressed |
| HAR_NO | SOF_NO | HAR_NO__SOF_NO | 4.44 | 135 | Compressed |
| BOL_IS | SKA_IS | BOL_IS__SKA_IS | 3.58 | 132 | Compressed |
| HAL_DK | ORU_SE | HAL_DK__ORU_SE | 5.21 | 130 | Compressed |
| BOK_NO | FLE_NO | BOK_NO__FLE_NO | 4.56 | 122 | Compressed |
| BOL_IS | BRE_IS | BOL_IS__BRE_IS | 4.56 | 103 | Compressed |
| BOK_NO | HAR_NO | BOK_NO__HAR_NO | 4.30 | 102 | Compressed |
| ALB_DK | SKA_DK | ALB_DK__SKA_DK | 5.97 | 91 | Compressed |
| ALB_DK | HAL_DK | ALB_DK__HAL_DK | 4.56 | 69 | Compressed |
| ALB_DK | ORU_SE | ALB_DK__ORU_SE | 4.59 | 68 | Compressed |
| FAL_SE | HEL_DK | FAL_SE__HEL_DK | 11.51 | 48 | Compressed |
| BAK_IS | VOP_IS | BAK_IS__VOP_IS | 3.53 | 45 | Compressed |
| FAL_SE | SVAN_SE | FAL_SE__SVAN_SE | 7.05 | 34 | Compressed |
| HEL_DK | SVAN_SE | HEL_DK__SVAN_SE | 7.83 | 17 | Compressed |
